# Deconstructing Gastrulation at the Single Cell Level

**DOI:** 10.1101/2021.09.16.460711

**Authors:** Tomer Stern, Sebastian J. Streichan, Stanislav Y. Shvartsman, Eric F. Wieschaus

## Abstract

Gastrulation movements in all animal embryos start with regulated deformations of patterned epithelial sheets. Current studies of gastrulation use a wide range of model organisms and emphasize either large-scale tissue processes or dynamics of individual cells and cell groups ^1,2,11–13,3–10^. Here we take a step towards bridging these complementary strategies and deconstruct early stages of gastrulation in the entire Drosophila embryo, where transcriptional patterns in the blastoderm give rise to region-specific cell behaviors. Our approach relies on an integrated computational framework for cell segmentation and tracking and on efficient algorithms for event detection. Our results reveal how thousands of cell shape changes, divisions, and intercalations drive large-scale deformations of the patterned blastoderm, setting the stage for systems-level dissection of a pivotal step in animal development.

## Results and discussion

During the first two hours of development, the *Drosophila* embryo goes through 13 synchronous cleavages and arranges most of its nuclei in a syncytial monolayer under the common plasma membrane. The next hour is characterized by cell cycle arrest and zygotic genome activation, which transforms the nuclear syncytium into a patterned epithelial sheet. The newly formed and elaborately patterned epithelium then proceeds to gastrulation. As in embryos of other animals, *Drosophila* gastrulation relies on spatial and temporal control of cell divisions, cell shape changes, and cell intercalations. Each of these behaviors has been associated with distinct aspects of gastrulation ^14–17^ and has been an object of intense studies using genetic, cell biological, and more recently, biophysical approaches ^1,18–26^. Most of these studies, however, focus either on cellular processes driving gastrulation or on large-scale tissue deformations. Recent advances in microscopy and image processing create a unique opportunity for integrating these complementary viewpoints ^9–12^. Our study capitalizes on these advances to provide the first complete view of gastrulation at the single-cell level.

Our strategy for deconstructing gastrulation has two main ingredients. The first ingredient is a custom-made approach to cell segmentation and tracking (Figure 1A and [supplementary videos 1](https://www.youtube.com/watch?v=kC11Upr30JY),2,3). This process starts by forming a two-dimensional (2D) Mercator projection of the apical side of the epithelium sheet for each time point ^27^. Next, the cells in the projection of the first time point are segmented and assigned unique track ID ^28^. Cell boundaries and track IDs are then propagated iteratively and automatically over time. This is done by first deforming each projected image to accurately match the position of each cell with the same cell at the next time point, using Maxwell’s Demons algorithm ^29^. Once cell positions are matched, cell boundaries and track IDs are copied to the next time point, and then fine-tuned using the watershed transformation ^30^. Lastly, the segmented and tracked 2D projection at each time point is used to generate a 3D polygonal mesh that reveals the apical surfaces of the cells and provides an in-toto map of cell-cell adjacencies ^31^. We optimized each step of this pipeline, enabling it to carry out ∼1 million reliable single cell segmentations in a typical light sheet imaging dataset from a gastrulating embryo (Figure S1A-F).

**Figure 1:**
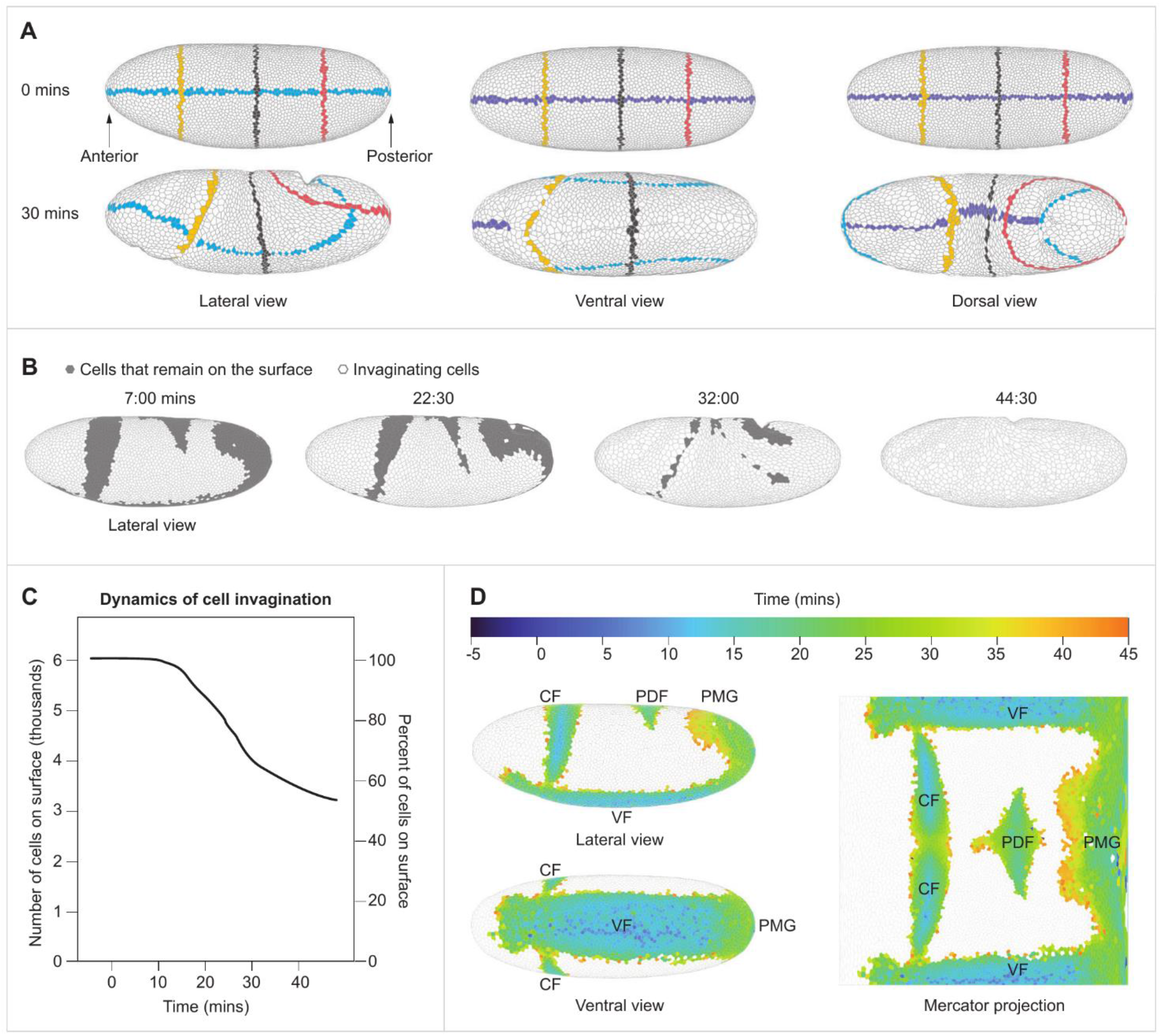
Cell segmentation and tracking, and cell invagination mapping. (A) Demonstration of whole embryo single cell segmentation and tracking. Lateral, ventral, and dorsal views of the 3D polygonal mesh at 0 and 30 minutes. Cells intersecting with the mid-coronal plane, the mid-sagittal plane and three transverse planes have been distinctly colored to demonstrate cell tracking and to reveal how the embryo deforms over time. (B) Lateral view of the embryo at four time points showing invaginating cells (gray) and cells that remain on the surface (white). (C) Quantification of the number and percent of cells (left and right vertical axes, respectively) remaining on the surface of the embryo over time. (D) Left: Projection of invagination sites during the first 45 minutes of gastrulation over the 3D blastoderm shell in lateral and ventral views. Right: Mercator projection of the invagination sites. Each cell is color-coded for the first time point in which it is no longer visible from the outside of the embryo. *t*=0 was set as the beginning of apical constriction in the invaginating mesodermal cells. CF – Cephalic Furrow, VF – Ventral Furrow, PDF – Posterior Dorsal Fold, PMG – Posterior Midgut.

Automated detection of dynamic behaviors of single cells and cell groups is the second key ingredient of our computational strategy for deconstructing gastrulation. Given accurate information of cell volumes and areas, detection of single cell behaviors (division, internalization, and columnar-to-squamous transition) is relatively straightforward. Large scale detection of events involving cell groups, such as localized cell intercalations, is more demanding, both algorithmically and computationally. We used our recently published algorithm for searching dynamic cell behaviors, which is based on optimized graph exploration and multivariable time series matching ^32^. To provide an idea for the scale of detected events, in one of the analyzed live imaging datasets from an embryo that started with 6085 cells in the blastoderm, we detected 2808 cell invaginations, out of which 1153 were in the ventral furrow, 582 in the cephalic furrow, 728 in the posterior midgut, and 345 in the posterior dorsal fold. In addition, we detected 1110 cell divisions, 1881 cell intercalations, and 176 columnar-to-squamous cell shape changes.

We illustrate our computational approach by focusing on one of the most striking aspects of *Drosophila* gastrulation. Specifically, spatially patterned cell invaginations result in progressive loss of almost 50% of all cells originally present on the surface of the blastoderm (Figure 1B,C). Accurate tracking allows us to map the participating cells to the early blastoderm, thereby providing a fate map for cell invagination (Figure 1D). This type of retrospective mapping will also be essential for analyzing how cell behaviors in the remining regions of the embryo contribute to large scale tissue deformations and compensate for cell invagination through spatially controlled intercalations, shape changes, and divisions.

We start by focusing on the germ-band (GB), a domain of the embryo that converges and extends as a consequence of multiple cell intercalations. Most of these events involve cell quartets and proceed through the so-called T1-transition, in which two nonadjacent cells come together and split the “orthogonal” cell pair (Figure 2A) ^33^. Another intercalary event involves more than four cells, and proceeds through a configuration where several contracting interfaces generate a so-called rosette state ^34^. We mapped the two bilaterally symmetric blastoderm primordia for the GB using our event detection algorithm to identify all T1s and rosettes during the first 45 minutes of gastrulation (Figure 2B,C).

**Figure 2:**
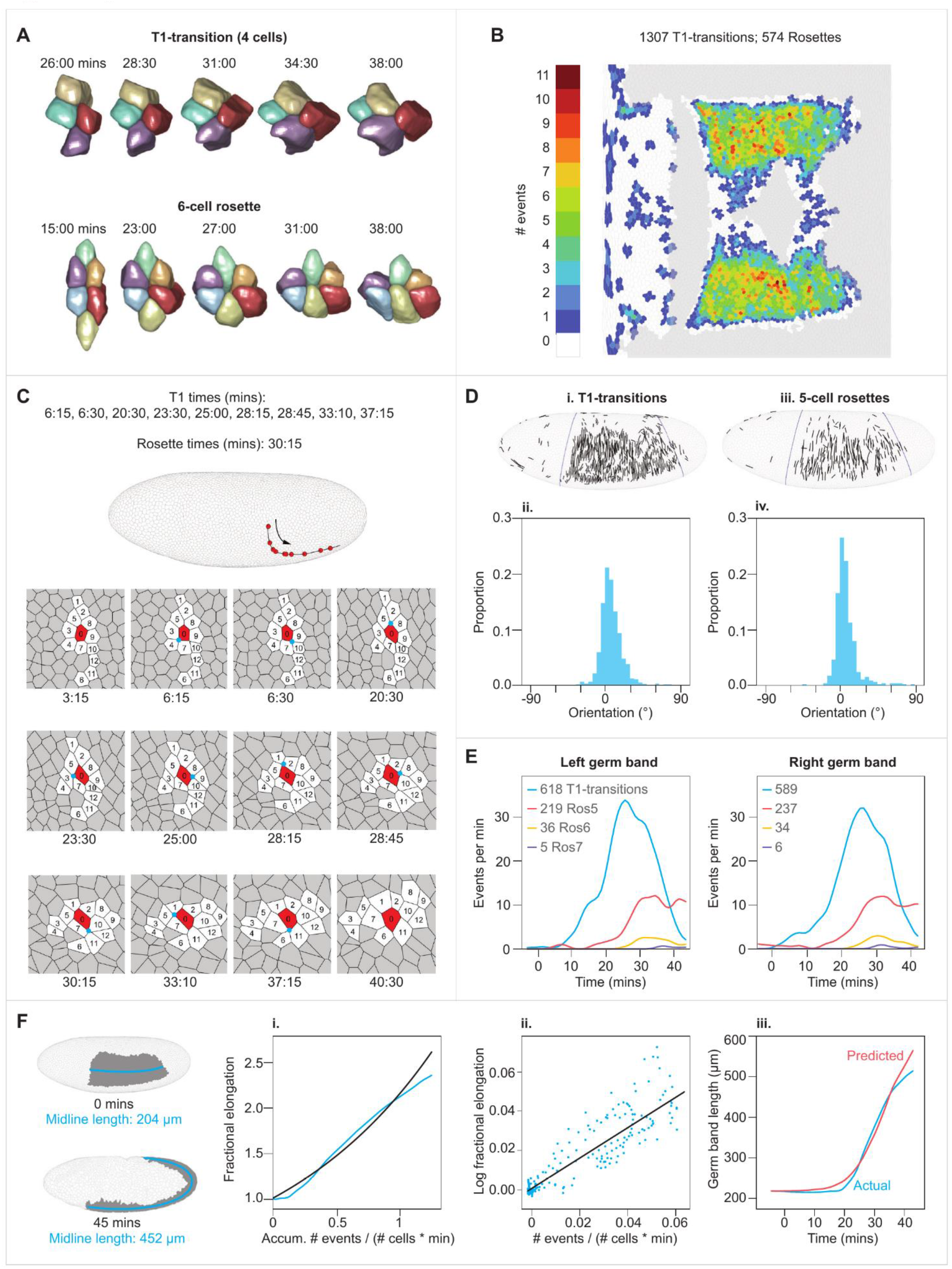
Dynamics of cell intercalations. (A) Three-dimensional reconstructions of a representative T1-transition and a 6-cell rosette. (B) Mercator projection showing the number of intercalary events in which a cell participates over the first 45 minutes of gastrulation (both T1s and rosettes). (C) Top: a lateral view of the embryo showing the trajectory of a single cell that participates in 10 sequential intercalation events. Each red dot indicates the position of the cell at the moment of an event. Bottom: segmented view of this cell with its neighbors. The central vertex at each intercalation is marked by a cyan circle, and numbers inside cells indicate cell identities. The first and last panels show the configuration of the tissue prior to and resulting from all intercalations, respectively. (D) Top: spatial mapping of intercalary events based on the original positions of converging cells in the blastoderm: T1-transition (i) and 5-cell rosette (iii). Orientations shown from the lateral view of the embryo. Bottom: histograms of the orientation angles of T1-transitions (ii; avg. ± S.D. = 7.7° ± 12.6°, *n*=1207) and 5-cell rosettes (iv; avg. ± S.D. = 7.3° ± 13.1°, *n*=456). (E) Comparison of the temporal frequencies of T1-transitions and 5-, 6-, and 7-cell rosettes between left and right sides of the embryo. (F) (i) Left: lateral view of the 3D surface of the embryo at 0 and 45 minutes, demonstrating convergence and extension of the germ band (gray). The length of the germ-band is measured as the length of the midline (cyan). Right: comparison between the actual fractional elongation of the germ-band plotted as a function of the normalized cumulative number of events (cyan), and that predicted by our mean-field model (black; see text). (ii) Linear regression of our mean-field model (black). Each dot (cyan) is the log fractional elongation of the germ-band over one minute as a function of the normalized cumulative number of events during the same time period. (iii) Parametric plot of the time dependent germ-band length constructed based on the mean-field model and measured number of intercalary events. Blue – empirical measurement, red – predicted.

Individual intercalary events can produce global changes in the shape of a primordium only if they share a common orientation. Oriented intercalations have been documented in small regions of the embryo ^35^, but previous studies could not track all events, especially as the extending GB rounds the posterior pole of the embryo. To examine orientational order, we mapped intercalary events back to blastoderm surface and represent their orientations as lines between the centers of cells that are brought into contact by an intercalation (Figures 2D and S1G). Although individual intercalations within the GB occur at different times and in different positions, all are strictly aligned with the overall convergence of the tissue and the dorsal ventral axis of the embryo. This global alignment argues strongly that directionality of intercalary events is set already at the blastoderm stage.

Two different cells within the GB can undergo very different numbers of intercalary events; the number can be as low as one and as high as eleven (avg. ± S.D. = 4.18 ± 2.15; Figure 2C). In spite of this high variability at the single cell level, the integrated number of events are very similar on the two sides of the same embryo (Figure 2E). This observation prompted us examine the dependence between the number of intercalations and global tissue elongation. By plotting the fractional tissue elongation versus the number of intercalations that had happened by time *t* we found a surprisingly simple dependence (Figure 2Fi). This behavior can be rationalized using a mean-field model whereby each neighbor exchange, a T1 transition or a rosette, has the same effect on tissue elongation, based on the fact that all events show directionality. Furthermore, we assume that effects happening in different time intervals are independent of each other. Such memoryless and additive model gives rise to an exponential dependence between the number of events and elongation: *L*(*t*)/*L*(0) = exp(*αE*(*t*)/*C*), where *E*(*t*) is the cumulative number of events by time t, *C* is the total number of cells in the primordium, and *α* is a proportionality factor. This model provides a direct connection between oriented cell intercalations and global deformation. Fitting the model to data (Figure 2Fii,iii; *α*=0.762; *R*^*2*^=0.758) reveals that in a tissue with *C* cells, 0.91*C* events would double its length. Future work will be needed to understand the close match between the number of events and the size of the tissue.

Despite significant intercalation of cells, the total area occupied by the GB increases only by 1.46-fold during the elongation process, an increase attributable to a slight increase in the surface area occupied by individual cells (see below; Figure S2A-D). To test whether cell proliferation might provide an alternate strategy compensating for cell loss due to invagination, we identified and followed all dividing cells during early gastrulation. In *Drosophila*, post-blastoderm mitoses can be assigned to 25 domains defined by patterned expression of a mitosis-promoting protein phosphatase ^15,36^. We constructed a spatiotemporal map of these mitoses, recovering all of the previously described mitotic domains, along with the intra-domain mitotic waves (Figures 3A and S3). Consistent with the early description^10^, during this period the vast majority (96%) occur outside the GB. Using our 3D segmentation and tracking pipeline, we then followed single-cell dynamics of all dividing cells. During the course of each division (Figure 3B), cells round up but conserve volume, such that the two daughter cells have a combined volume equal that of the parent cell (Figures 3Ci and S2E,F). Cell rounding does however cause a transient increase in apical area, coupled with a similarly transient apicobasal shortening (Figure 3Ciii,iv). Given the large number of cells dividing at any moment, these transient increases add up to a 2.43-fold increase in the area occupied by all dividing cells (Figure 3Cii) and provide a significant compensation for cell loss during gastrulation.

**Figure 3:**
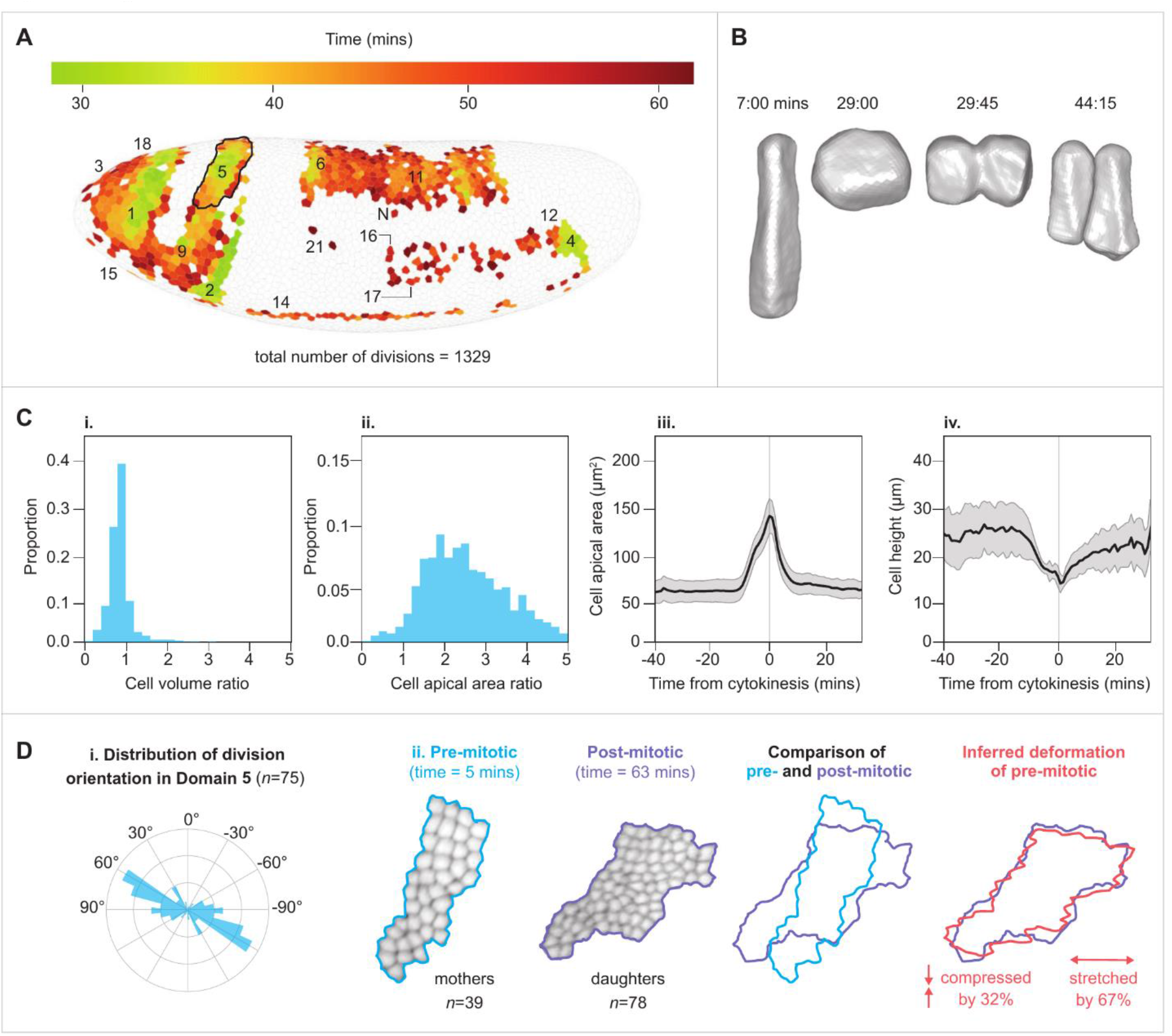
Dynamics of cell divisions. (A) A spatial map of cell divisions where dividing cells are shown in their original positions in the blastoderm stage, and color indicating their time of completion of their division. Mitotic domains indicated by the nomenclature proposed by Victoria Foe ^15^. (B) Three-dimensional reconstruction of a dividing cell, showing transient increase in apical area upon rounding up (*t*=29:00). (C) i-ii) Histograms of the ratios of final to initial cell volume (i) and final to initial cell apical area (ii) in dividing cells. In both histograms *t*_*final*_*=43:00*. For cell areas *t*_*initial*_*=0:00* and for cell volumes it is the time at which cellularization is completed in the entire embryo (*t*_*initial*_*=17:00*). iii-iv) Average and S.D. of apical area and apicobasal cell height from domain #1. Values represent the apical area of the mother cell and the summed apical areas of both daughter cells. The sharp increase in area toward t=0 is followed by decrease to approximately pre-division levels, demonstrating that cell divisions have a significant, yet transient contribution to tissue expansion. (D) Spatial organization of cell divisions in domain 5: (i) Polar histogram of the distribution of division orientations within the domain relative to the DV-AP axes of the embryo (90^0^ = dorsal). (ii) Shapes of the domain before and after the completion of divisions. To a good approximation, the two shapes are related to each other by a simple area preserving linear transformation, which includes a stretch and a compression factor.

We found that that several mitotic domains (1, 2, 5, 6, and 18) display highly aligned directions of cell divisions and are, as a consequence, stretched in the direction of common cell division and compressed in the orthogonal direction (Figures 3D and S1H). Such stretching/compressing reshaping is a direct consequence of the fact that, after the described overshoot, the apical area of an approximately hexagonal parent cell is distributed between two smaller adjacent hexagonal daughter cells. The postmitotic shapes of several domains are indeed consistent with this type of area-preserving transformation, showing that patterned epithelial domains can be reshaped not only by directed cell intercalations, but also by directed divisions. Similar polarized divisions have been previously reported for domain 4 ^37^. Another domain reshaping operation is based on anisotropic cell shape changes, even in the absence of cell intercalations and divisions (Figure 4A). This strategy is followed by the amnioserosa primordium, a narrow band of cells on the dorsal side of the embryo. In response to several locally expressed transcription factors and signaling molecules, cells within the amnioserosa primordium change their shapes from columnar to squamous, expanding their projected apical area by 3.10-fold, with minimal changes in cell volume (Figures 4B,C and S2 and [supplementary video 4](https://www.youtube.com/watch?v=TuKAU2KDiYQ)) ^16,38^.

**Figure 4:**
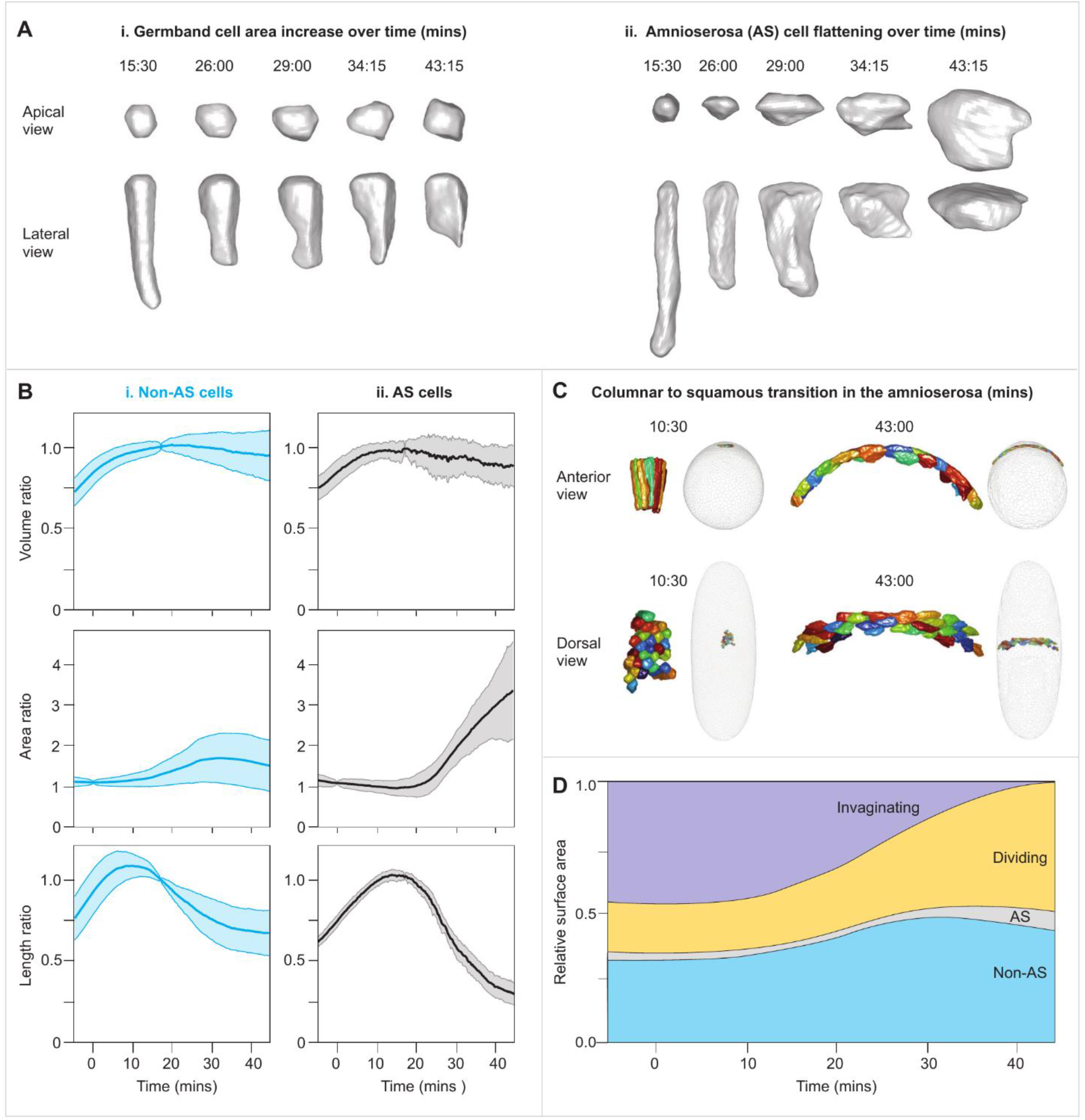
Dynamics of columnar to squamous cell shape transformation. (A) Three-dimensional reconstruction of the geometric transformations of cells from the germ-band (i) and from the amnioserosa (ii). (B) Dynamics of the volume ratios, area ratios and apico-basal length in AS and non-AS cells. Black and blue lines are averages, and shaded colors are the S.D.s over time. The completion of cellularization in the AS (approximately 11 minutes) and non-AS cells (approximately 8 minutes) is indicated by peak in the cell length. At that point, volumes remain relatively constant, and apical surface areas begin to increase. (C) Three-dimensional reconstruction of AS cells before and late in the columnar to squamous transition, shown as anterior (top) and dorsal (bottom) views. In the gray whole embryo views, the locations of the colored AS cells are shown viewed from the anterior end (upper images) or from the dorsal side (lower Images). (D) The fraction of embryo surface occupied by cells undergoing each of the four behaviors over time, demonstrating the compensation of area loss due to cell invagination.

Detection of all instances of cell intercalation, division, and columnar-to-squamous transition allowed us to determine how each of these small-scale events contribute to the large-scale areal expansion that compensates for the loss of invaginating cells (Figure 4D). Although the cells of amnioserosa undergo the largest relative expansion of their apical area, the fraction of the amnioserosa primordium that remains on the surface is small and contributes only 5% to covering the 46% area deficit. The majority of the deficit (30% out of 46%) is compensated by the dividing cells, which can increase their areas quite significantly during individual division events. The residual nondividing regions of the blastoderm, including the previously described germ band, undergoes an average 1.35-fold expansion of apical surface area, covering the remaining 11% of the area debt.

Our study demonstrates how recent advances in microscopy, image processing, and machine learning can be used to deconstruct gastrulation in the whole embryo into behaviors of individual cells and small cell groups. As a first illustration of this approach, we characterized the large-scale areal expansion that accompanies well-characterized processes of epithelial invagination and GB expansion. Each of the constituent behaviors is a complex regulatory module, reflecting the action of transcriptional, signaling, and cytoskeletal circuits. Our approach bridges the gap between molecular and cellular studies and biophysical models that view epithelium within the continuum mechanics framework where the main players are no longer individual cells, but coarse-grained fields of deformations and forces. Having access to cell behaviors throughout embryo promises to provide the much-needed boundary conditions and constitutive laws for these models. The early *Drosophila* embryo is ideally positioned for making this connection, but we expect that other developmental models will soon follow suit.

## STAR Methods

### Detection of cell intercalation

T1-transitions (a.k.a 4-cell rosettes) were detected according to our previously developed high accuracy algorithm for template-based mapping of dynamic motifs in tissue morphogenesis ^32^. For high order rosettes we updated our algorithm so that a rosette must meet all following criteria:

#### Topological criteria

- Every cell from the rosette has at least two neighbors from within the rosette.
- Every cell from the rosette must have at least one neighbor that is not from the rosette.
- There is no connected component of cells that do not belong to the rosette and that is surrounded only by cells from the rosette.
- There is no cell from the rosette whose removal from the rosette disconnects two other cells from the rosette (i.e. the removal of that cell leaves no path between two other cells in the rosette that passes only through the remaining cells of the rosette).
- Remove intercalations that cross invaginations: during invagination, cells from the two sides of a furrow (e.g. the ventral furrow) may appear in the polygonal mesh as undergoing intercalation, although biologically this is not the case. To identify and discard these intercalations, assume an intercalation event at time point *t* = *T* that includes *n* cells. For each pairs of cells (*c*_*i*_, *c*_*j*_) find the all the cells along the shortest path from *c*_*i*_ to *c*_*j*_ at time *t* = 0. If any of the cells along this path will invaginate before time point *T*, we conclude that the intercalation is a result of invagination rather than active intercalation.

#### Geometric criteria

- The maximal apical area of a cell from a rosette is no more than 10 times the minimal apical area.
- The maximal pairwise Euclidean distance between the boundaries of a pair of cells from the rosette is (*n*− 2) microns, where *n* is the number of cells within the rosette.
- We define an internal interface to be an interface of the rosette if all the cells that meet at its two vertices belong to the rosette. An external interface is an interface between two cells of the rosette that is not an internal interface. We require that the sum of lengths of all internal interfaces in a rosette will be larger than one third the median length of external interfaces.

#### Overlap and duplications criteria

- If the cells of an intercalary event “A” are a subset of another intercalary event “B”, and the time difference between the two events is no more than 5 minutes, we remove event “A”.
- If a group of *n* cells meet all described criteria at more than one time point, we keep only the event at the time point in which the maximal pairwise Euclidean distance between cell boundaries is minimal.
- We remove intercalations that include a dividing cell. Since a dividing cell can exert significant mechanical force onto its neighbors and lead to cell intercalation, we remove any intercalation in which one of the cells is dividing in the interval between 2 minutes before and 8 minutes after the intercalation event.

Lastly, since our previous benchmark for the algorithm was done only on simulated data and on the extending germ, all detected intercalations outside of the germ-bands were validated manually.

### Calculation of intercalation orientation with respect to the ventral midline: (Figure S1G)

A. **Generation of the ventral midline:** We define the ventral midline as the line on the 3D surface of the embryo that passes in the middle of the domain of cells that will later invaginate as a part of the ventral furrow. To accurately mark the ventral midline, we rely on the 2D Mercator projection of cell invagination map (Figure 1D in the main body of the manuscript), by calculating a distance transform for pixels in the invaginating regions to reach pixels in the non-invaginating regions. The distance transform highlights the ridge of pixels that are farthest from the non-invaginating parts of the embryo, which the user marks with a straight line. Lastly, we transform the points on the marked line back to the original 3D embedding space of the embryo, which will serve us as the ventral midline.
B. **Calculation of the DV orientation at a point on the blastoderm surface:** Given a point (*x*_*vm*_, *y*_*vm*_, *z*_*vm*_) on the ventral midline, we construct a plane *P* that is orthogonal to the ventral midline at that point. The intersection of *P* with the surface of the blastoderm has a ring-like shape which corresponds to a cross-section of the embryo. The DV orientation at a point (*x*_*IL*_, *y*_*IL*_, *z*_*IL*_) on the intersection line is the orientation of the tangent to the intersection line at that point.
C. **Calculation of the orientation of an intercalation event:** The blastoderm projection of a T1-transition event in which the cells *c*_*A*_ and *c*_*P*_ are at the two ends of the contracting interface, is defined as the line between the centroids of cells *c*_*A*_ and *c*_*P*_: (*x*_*A*_, *y*_*A*_, *z*_*A*_) and (*x*_*P*_, *y*_*P*_, *z*_*P*_), respectively, at time point t=0. The orientation of this line is the angle between itself and the DV orientation at its middle: 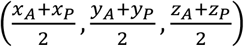. In the case of a rosette (5 cells or more), cells *c*_*A*_ and *c*_*P*_ will be the two cells that the distance between their centroids is the highest among all cell pairs from the rosette, at time point t=0.

### Labeling of mitotic domains

(Figure S3) Identification of the set of cells belonging to each mitotic domain was done manually by comparison of the anatomical distribution and temporal dynamics shown in our cell division maps (Figure 3A) to these reported by Foe ^15^. Domains 3, 8, 15, 18 and 19 did not manifest clear characteristics and were therefore marked with limited confidence. Otherwise, the temporal and geometric characteristics of the identified domains were in strong agreement with Foe.

### Projection of division orientation onto the early blastoderm

(Figure S1H) Motivation: the goal of projecting the orientation line onto the blastoderm is to reverse changes in orientation that have occurred due to tissue deformations throughout gastrulation. For instance, assume a mitotic domain that is shaped as a rectangle, and all cells within the domain are dividing along the short axis of the domain. If the domain does not change its position over time, then all division orientation lines would appear collinear, in which case our method will not be needed. However, if the domain itself is being rotated by 90 degrees over time, as some domains do, then the division lines would distribute over 90 degrees, which would obscure the fact that all cells share the same orientation with respect to the mitotic domain. Method: given a cell division event wherein the two daughter cells appear for the first time at time point *t*, we first calculate the orientation line of the division as the line between the cell centroids of the two daughter cells. To correct for local deformations, we first identify the set of *N* first and second neighbors of the mother cell at time point *t* = 0 (i.e. all cells whose shortest path to the mother cell at time point 0 is ≤ 2). Then, we calculate the (*x, y, z*) centroid coordinates of each of the *N* neighboring cells, giving us the matrix: *C*_(0)_ ∈ ℝ^*N*×3^. We continue by generating the corresponding matrix of centroids for the same set of neighbors at time point *t*: *C*_(*t*)_. In case any of the *N* neighbors have divided prior to time point *t*, we calculate its centroid at time point *t* as the average between the centroids of the two daughter cells. Next, we fit an affine transformation *A* ∈ ℝ^4×4^ that optimizes the match of *C*_(*t*)_ to *C*_(0)_ (this requires turning *C*_(*t*)_ and *C*_(0)_ into homogenous coordinates by padding each of them on the right with 1^*N*×1^). Lastly, we apply *A* on the coordinates of the end points of the orientation line, which provides us an estimation of the orientation prior to tissue deformations. For visual clarity, we fine tune the position of the projected line by aligning its mid-point with the centroid of the mother cell at time 0, and update its length to be 5 microns so when plotted it would not cross the boundaries of the mother cell.

### Circular statistics of division orientation

Projecting each division line onto the blastoderm allows us to quantify the distribution of orientations with respect to the AP and DV axes of the embryo. As mitotic domains typically include not more than several tens of cells and are concentrated in a small region in the embryo, we can approximate these orientations based on the representation of the lines in the *ImSAnE* pullbacks (the 2D Mercator projection of the embryo surface). In this representation the vertical and horizontal axes of the image approximate the AP and DV axes of the embryo and allow immediate extraction of the orientations. Since the cells within the same domain at the two sides of the embryo are mirror-image of one another, we begin by manually marking the right half of the embryo and mirroring each division line from the right side to the left. Then, we calculate the orientation of each blastoderm projected division line as: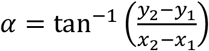, where (*x*_1_, *y*_1_) and (*x*_2_, *y*_2_) are the endpoints of the projected line. Since orientations are radially symmetric, and the symmetry in our case is synonymous, prior to calculating the histogram we double the number of measured orientations in each mitotic domain by adding for each angle α its opposing angle: α+180^0^, making the distribution diametrically bimodal. Polar distribution was then calculated with bins separated at values: (−5^0^, 5^0^, 15^0^, …, −5^0^).

The average orientation of divisions in a mitotic domain with angles {*α*_*j*_} was calculated as:

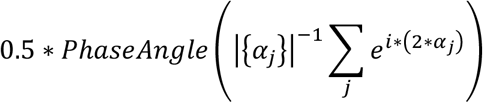

Note that in diametrically bimodal data, calculating the average directly on the set of angle results erroneously in an average that is orthogonal to the true average. To overcome this, we apply the angle doubling procedure (see 2 * *α*_*j*_ in the exponent and the 0.5 rescaling factor). The standard deviation of the angle set {*α*_*j*_} was calculated as: 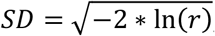, where *r* is the length of the mean resultant vector.

### Quantification of cell geometry

(Figure S2A-C,E) Cell volume at a given time point was calculated from the image of 3D segmentation as the total number of voxels assigned with the index of that cell × voxel volume (0.2619 *µm^3^*). Cell apical area was calculated based on the segmented triangle mesh (i.e., prior to its conversion to a polygonal mesh) by summing the areas of triangles assigned with the index of that cell. To calculate cell length along the apico-basal axis we first calculated an apico-basal (AB) axis. We define the AB axis as the line that is normal to the polygon representation of the cell in the 2D segmentation and passes through the centroid of the polygon. Then, we projected the 3D coordinates of all voxels of the cell from the 3D segmentation onto the normal line. The AB length of the cell was then calculated as the Euclidean distance between the projected voxels at the 1^st^ and 99^th^ percentiles along the line. Analyses of volumes and lengths were validated manually.

### Quantification of final to initial cell geometric ratios

(Figure S2D,F) For the calculation of ratio between final and initial cell apical area, apico-basal length, and volume, we used t=45 minutes as the final time point since this is when invaginated cells start coming back out. For apical area ratio we used the last time point before gastrulation movements begin for the initial value (t=0 minutes). For volume and length ratios we used the time at which cellularization over the entire embryo completes as the first time point (t=17 minutes).

### Calculation of germ-band length dynamics

Our strategy is to identify in the blastoderm a line of cells that spans the entire AP axis of the germ-band and is located at mid DV length (see Figure 2F left panel in the main text). The length of the germ-band at any time point throughout the development of this embryo is defined as the arc-length of a smoothed line that passes in proximity to the centroids of these cells. Our empirically validated assumption is that the identified line of cells will span along the entire germ-band at its mid-height at all later time points as well, which allows the calculation of length to be carried automatically from the second time point onward. Method: first, the blastoderm projected map of the number of intercalations per cell is presented, on which the user marks the cells passing in the middle of the germ-band (Figure 2B in the main text). Although any line that passes through the centroids of the midline cells will approximate the midline of the germ-band, it will most likely zigzag between the centroids of neighboring cells and therefore overestimate the length of the germ-band. Instead, we fit a three-dimensional smoothing spline curve through the centroids. Lastly, we estimate the length of the spline using linear fragmentation.

### Regression analysis

The goal of our model is to connect the relative elongation of the germ-band over a given time period with the density of intercalary events occurring over the same period: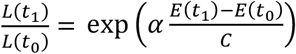, where *L*(*t*) and *E*(*t*) are the length of the tissue and the accumulated number of events at time *t, C* is the total number of cells in the primordium, and *α* is a dimensionless proportionality factor. Notice that *α* can be estimated using linear regression by applying a ln(*x*) function on both sides: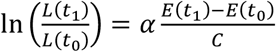. To generate sample data, we manually marked the right and the left germ-bands in two selected embryos based on the Mercator projection showing the number of intercalations per cell (Figure 2B in the main text), resulting in four quantified germ-bands. In each germ-band the fractional elongation of the tissue as well as intercalation density (# intercalations / # cells within the germ-band) were calculated in one-minute intervals, providing a total of 172 data points. Regression was then done using MatLab’s “regress” function.

### Software

All parts of the pipeline except for the implementation of the algorithm for generating subgraph induced connected components of size k, were developed in *MatLab* 2020b. For basic image processing tasks such as rotation, ROI cropping and format conversion we used *Fiji* [62]. Image segmentation and object classification in 2D for the first time point in each embryo were done using *Ilastik* [58]. Implementation of the algorithm for generating subgraph induced connected components of size k was done in c language by Shant Karakash and Berthe Choueiry (University of Nebraska-Lincoln, NE, USA) and used under their permission. Unrolling of the apical surface of the embryo was done using *ImSAnE* ^27^. 3D segmentation and tracking of cells relied on the deep-learning algorithm *CellPose* ^31^. Calculations of circular statistics relied on the circular statistics toolbox ^39^.

### Hardware

Code for the described pipeline was developed on a Dell PowerEdge R930 server carrying four Intel(R) Xeon(R) CPU E7-4850 v4 @ 2.10GHz with 16 cores each and 2TB RAM, with RedHat OS. Cell and nucleus segmentation and tracking, and all the analyses of the extracted motifs were done on Intel(R) Xeon(R) CPU E5-1620 v4 @ 3.5GHz, with 32GB memory and Windows 10 64 bits OS. 3D segmentation using the algorithm *CellPose* was done on a Nvidia P100 GPU with 16 GB memory.

### Live imaging

Light-sheet movies were acquired and reconstructed as described in ^1^. Embryos were imaged at rate ranging from frame per 15 to per 75 seconds. In all movies reconstruction resulted in an isotropic voxel dimensions of 0.2619 microns.

### Fly stocks

Embryos to be imaged were obtained from Gap43::mCherry/TM3 females or from Tub67a < CAAX – mCherry < sqh 3’ UTR. attp 2. / TM3, Sb females.

## Resource availability

### Lead contact

Further information and requests for resources should be directed to and will be fulfilled by the Lead Contact, Stanislav Y. Shvartsman.

### Materials availability

This study did not generate new unique reagents.

### Data and code availability

Raw image data and code are available upon request.

## Acknowledgements

We are grateful to Lucy Reading-Ikkanda for graphic design of the figures, to Lisa Brown (Simons Foundation) for fruitful discussions on image processing, to Trudi Schüpbach, Matej Krajnc (Jožef Stefan Institute), David Denberg, Tal Galili, and Yosi Keller (Bar-Ilan University) for fruitful discussions, to Matthew Cahn (Princeton University) and the IT of Simons Foundation for technical support with computing resources, to Reba Samantha and Laisa Eimont for bureaucratic assistance and to Heping Jiang for stock maintenance, and lastly to the Wieschaus Shvartsman lab members for continuous support.

