## Supplementary materials for "Deconstructing Gastrulation at the Single Cell Level"

### Supplementary videos

#### Video S1.

https://youtu.be/kC11Upr30JY

Development of an embryo expressing a membrane marker presented from four views (left to right): Dorsal, Ventral, Lateral-Right and Lateral-Left, related to Figure 1. Each frame was generated by extracting the surface located 5 microns beneath the outer surface of the embryo directly from the raw 3D light-sheet data. The movie spans from 5 minutes before to 45 minutes after the beginning of ventral furrow formation with temporal resolution of 1 frame every 15 seconds and displayed at a frame rate of 14 frames per second.

#### Video S2.

https://youtu.be/lKKNxIH3eCI

Results of our segmentation algorithm applied on the embryo shown in Video S1, related to Figure 1. The segmentation of a single time point generates a polygonal mesh, in which polygons correspond to cells.

#### Video S3.

https://youtu.be/TrBgukVDRV8

Results of our tracking algorithm applied on the embryo shown in Video S1, related to Figure 1. Polygons were color-coded randomly for the identities of the cells, allowing to track the cells visually over time and estimate the quality of the algorithm. In the event of cell-division each of the two daughter cells will be assigned a random color that is independent of the color of the mother cell.

#### Video S4.

https://youtu.be/TuKAU2KDiYQ

3D reconstruction of amnioserosa cells from an anterior view, related to Figure 4C. The movie spans from 5 minutes before to 45 minutes after the beginning of ventral furrow formation, with temporal resolution of 1 frame every 15 seconds and displayed at a frame rate of 28 frames per second. The movie begins halfway through the cellularization of these cells. Immediately after cellularization is completed columnar to squamous transition initiates, together with the lateral spreading of the cells, increasing the cross-sectional angular span of the domain from 20° to 115° within less than 30 minutes.

### S1. A pipeline for cell segmentation, tracking, division detection and polygonization

**Input:** A series of $T \in \mathbb{N}$ 3D volumes of an entire fly embryo, reconstructed at isotropic voxel size of *dxyz=0.2619* and with temporal intervals of *dt* seconds (between 15 and 75 seconds).

**Output:** A series of $T$ polygonal meshes, wherein the polygons of mesh $1 \leq t \leq T$ accurately overlap with the cell membranes in the input volume at time $t$, approximately 5 microns underneath the apical end of the cells. Two polygons at two different time points have the same polygon index if and only if they represent the same biological cell.

**Detection of apical surface (S1.A):** We begin by extracting from each 3D volume a 2D surface that is located 5 microns beneath the apical tip of the cells. We selected the -5 microns plane since it presented the highest contrast of the membrane marker. To extract this plane, we use the "*Autocontext*" module from the freely available image processing software *Ilastik* [1] to train a binary classifier that accurately separates the embryo (labeled "1"; including cells and yolk) from the background (labeled "0"). Lastly, we use morphological erosion by 19 pixels (~4.98 microns) to bring the surface of the binary mask into alignment with the -5 microns surface of the embryo.

**Unfolding of apical surface into four 2D mages (S1.B):** Although the apical surface of the cells is essentially a 3D object, we preferred to convert it first into a standard 2D image, as more segmentation and tracking tools are available for pixel-based data. To do so, we used the *ImSAnE* algorithm [2]. Given a 2D surface embedded within a 3D Euclidean space, *ImSAnE* divides the surface into four anatomically overlapping regions: Anterior, Posterior, Dorsal and Ventral. Then, it defines a coordinate system covering the surface of interest (SOI). The intensity value at each point on the SOI can then be mapped and displayed in the reference of the coordinate system, resulting in a 2D presentation of the fluorescence data. We will use the following notation for the transformation of a pixel from the original 3D embedding space into the flattened space: $TF_{anterior}(x, y, z) = (u, v)$. Notably, *ImSAnE* also provides us with the inverse of the transformation: $TF_{anterior}^{-1}(u, v) = (x, y, z)$. We use *ImSAnE* to transform the surface defined by the boundaries of the binary object we described in the previous section into four overlapping 2D images, to which we will refer as "pullbacks".

**Segmentation of first time point (S1.C):** Our strategy for segmentation and tracking is through watershed seed propagation [3]. In our data, this strategy begins by segmenting the four pullbacks of the first time point, based solely on the pixel content of these images. This is followed by iteratively propagating the segmented cells to the next time points using non-rigid image registration [3]. To segment the pullbacks of the first time point we used the *Autocontext* module in *Ilastik*, which calculates the probability of each pixel to represent a cell membrane, as the basis for seeded watershed segmentation (pixel connectivity = 4). During this process, each cell (i.e., a connected component of pixels) at each pullback is assigned a unique index from 1 to the number of cells within that pullback.

**Assignment of coherent cell indices:** Proper cell tracking requires that each biological cell will have a unique and distinct cell index throughout the entire time-lapse. Arbitrary cell indexing, as described in the previous section, is guaranteed not to meet the required criterion. For instance, if a cell located in the overlap region of the dorsal and the ventral pullbacks, it is unlikely that by mere chance it will be given the same cell index. Therefore, we identify all the appearances of each of the ~6000 biological cells in all four pullbacks and assign all appearances the same unique cell index. The decision of whether two cells in two different pullbacks represent the same biological cell is done based on the level of their distance in the original 3D embedding space (i.e., before flattening the apical surface).

**Segmentation propagation and tracking:** Having the four pullbacks of the first time point properly segmented and indexed, we are now ready to describe the propagation step of the segmentation from time point $t$ to $t+1$. At this point we have the four raw (unsegmented) pullbacks at time points $t$ and $t+1$, and the segmentations of the four pullbacks from time point $t$. First, we apply the non-rigid registration method *Demon's algorithm* [4] to estimate the displacement field of each of the four pullbacks at time $t$ to the corresponding pullback at $t+1$. To this end, we used *MatLab's* implementation - *imregdemons*, using 8 pyramid levels, 1000 iterations per level, accumulated field smoothing of 2.5, and an initial subsampling of both source and target images by a factor of 2 in both X and Y axes. We use the calculated displacement field $D \in \mathbb{R}^{M*N*2}$, which displaces a pullback of size $M*N$ at time point $t$ to that at time point $t+1$, to propagate the segmentation of this pullback using nearest neighbor interpolation. This provides us an approximation to the segmentation of the pullback at time point $t+1$. To refine the propagated segmentation, we first apply morphological erosion to remove the three outer layers of pixels from each cell, then use the remaining pixels as seeds for watershed segmentation. Lastly, to guarantee that the indexing of the cells has not changed as a result of the applied watershed, we update the index of each resulting cell to that of its seed.

**Division detection (S1.D):** The watershed seed propagation approach provides highly accurate cell segmentation for live movies of membrane labeled tissues, and it has the advantage of tracking the cells at the same time. However, it has the inherent limitation of being unable to detect the presence of a new cells not propagated from the previous time point, such as the two daughter cells following a division event. To overcome this limitation, we integrated into our platform the deep learning segmentation algorithm "*CellPose*" [5]. *CellPose* has the advantage of segmenting each image independently and can therefore identify the two daughter cells following division. To use *CellPose* for division detection, we follow the registration-based segmentation of the pullback with a segmentation using *CellPose*. Then, for each cell segmented by the watershed seed propagation approach ($C_{ws}$) we declare it as a cell division event, if there are two segmented cells in the *CellPose* segmentation ($C_{CP1}, C_{CP2}$), such that:

1. $Area(C_{CP1}) \geq 35\ \mu\mathrm{m}^2$ and also $Area(C_{CP2}) \geq 35\ \mu\mathrm{m}^2$.
2. $0.5 \leq \frac{Area(C_{CP1})}{Area(C_{CP2})} \leq 2$
3. At least 85% of the area of $C_{ws}$ is covered by $C_{CP1}$ and $C_{CP2}$.
4. At least 85% of the area of each of $C_{CP1}$ and $C_{CP2}$ is covered by $C_{ws}$.
5. $\frac{OuterIntensity + InnerIntensity}{2} \leq InterfaceIntensity$, where (see Figure S1D):
    a. $OuterIntensity$ is the median pixel intensity at the non-interfacing membranes.
    b. $InnerIntensity$ is the 5th percentile intensity inside the two cells.
    c. $InterfaceIntensity$ is the median pixel intensity at the interfacing membranes.

Once all criteria are met, we replace $C_{ws}$ with $C_{CP1}$ and $C_{CP2}$ and assign each daughter cell a new and unique cell index.

So far, our pipeline results in the segmentation, tracking and detection of divisions of the entire time-lapse, where each time point is represented as four pullbacks. Next, we will describe how we reintegrate the four pullbacks at each time point back into the initial 3D configuration.

**Generation of 3D triangle mesh and projection of 2D segmented cells onto it (S1.E):** We begin by generating a triangle mesh at -5 microns beneath the apical tip of the cells. Let $TriMesh = (F, V)$ be the triangle mesh, where $F = \{(i,j,k)\}, 1 \leq i \neq j \neq k \leq |V|$ is the set of faces, which contains all triplets of

vertex indices composing each face, and $V = \{(x_i, y_i, z_i)\}, (x_i, y_i, z_i) \in \mathbb{R}^3$ is the set of vertices, which contains all 3D coordinates of the vertices composing the triangles. This was done by smoothing the binary mask (generated in the beginning of the pipeline) through convolution with a 3D gaussian kernel that has a standard deviation of 3 voxels, followed by calculating the iso-surface of the smoothed mask at iso-value 0.5. Our goal now is to index each triangle face $f = (i, j, k) \in F$ according to the cell to which it belongs, based on the segmentations of the pullbacks. Assume that $f$ is located at the anterior pole of the embryo. The geometric centroid of $f$ is the average coordinate of its three vertices. We define the index of $f$ to be the index assigned to the pixel $p = (u, v)$ from the segmentation of a pullback which minimizes: $\left|f_{centroid} - TF_{anterior}^{-1}(u, v)\right|_2$. Simply put, we search for the pixel within the pullback, which has the minimal Euclidean distance to $f$ within the original 3D (embedding) space. For instance, if $f$ is located at the tip of the anterior pole of the embryo, then the matching pixel will be the pixel located in the exact center of the anterior pullback.
As long as the triangles in the triangulation are sufficiently small, this procedure is valid by itself. However, a face located at the longitudinal center of the embryo can potentially be matched with a pixel from any of the four pullbacks, due to the significant overlap between the pullbacks. If the quality of the segmentation at a given position within any of the pullbacks is equal, it makes no difference to which pullback the nearest pixel belongs. However, from a mathematical perspective, pullbacks are bound to present some spatial distortions which reduce the quality of segmentation. For instance, the anterior pullback appears less distorted at the center of the image, which corresponds to the apical pole of the embryo, and more distorted the farther we are from the center. Therefore, we prefer to limit the matching of $f$ to the pullback that is least distorted in the vicinity of $f$, and is therefore likely to have the most accurate segmentation there. To identify that pullback, we search the nearest pixel to $f$ according to the following criteria:

1. If $f_{centroid}$ is in the 20% anterior most region of the embryo, search only in the anterior pullback.
2. If $f_{centroid}$ is in the 20% posterior most region of the embryo, search only in the posterior pullback.
3. If $f_{centroid}$ is within the 20-80% along the AP axis and in the dorsal half of the embryo, search only in the dorsal pullback.
4. Otherwise, search only in the ventral pullback.

**From triangle mesh to polygonal mesh (S1.E):** At this point, each of the several thousand cells at each time point is represented by an average of several hundreds of mesh triangles, which would lead to high time and memory consumption in downstream analyses. Therefore, our last step is to convert the triangle mesh into a polygonal mesh, wherein each cell is represented by a single polygon, and each interface between neighboring cells will be represented by a single edge. The set of vertices of cell *c* in the polygonal mesh is the subset of vertices of cell *c* in the triangle mesh which are in contact with at least two additional cells (i.e., they are at the meeting point of three cells or more). In addition, an edge exists between two vertices in the polygonal mesh, if and only if both vertices are in contact with at least two mutual cells.

**3D Segmentation and tracking of external cells (S1.F):** Having the apical surface of the cells segmented and tracked allows us to segment and track the cells in three-dimensions with minimal efforts. To this end, we used *CellPose* to segment the raw 3D volumes. Improving the performances of the neural network used by *CellPose* on our images requires training it on examples of accurately segmented 2D images from our dataset (Note: *CellPose* has the advantage of using 2D examples to train segmentation of both 2D and 3D data). Fortunately, the generation of such examples close to the apical end of the cells is already an inherent part of our pipeline for 2D segmentation, thereby requiring preparation of 2D examples only close to the basal end, where imaging contrast is inferior due to unavoidable scattering. Once the training of the network and the segmentation of all volumes is completed, we are left with the task of tracking the segmented cells over time. To this end, we rely on the fact that at a given time point, the 2D polygon of a cell from the 2D segmentation is entirely, or mostly, contained within the volume of the same cell in the 3D segmentation. To update the indices of all 3D segmented cells, we follow the following routine: for each 2D polygon, identify the cell in the 3D image that contains the centroid of the polygon, and update the index of the 3D cell to the index of the polygon. In case a 3D cell is not reindexed throughout this procedure (i.e., it is not in overlap with the centroid of any polygon), or is reindexed multiple times (i.e., it is in overlap with centroids of multiple polygons), we remove it from the image by zeroing all its voxels.

**Method limitations:** Our mechanism for detecting an event of cell division is based on a hard-coded set of parameterized thresholds. On our data, we estimate the false positive rate of this approach to be within the range of 5-10%, and of false negative rate as 0-10%. Reducing the error level to 0% requires within the order of 2-3 hours of manual curation per time lapse. While we find the required manual efforts to be well within reason, given the intense labor associated with generating a single high-quality 3D time lapse at cell resolution, streamlining the generation of processed data may require a more adaptive approach with tolerable error rates.

The second and main limitation of our method stems from its inability to identify new cells appearing on the surface of the embryo. To detect cell division events, we integrated the *CellPose* into our pipeline and relied on a series of clear visual criteria as further validation. However, in addition to cell division, cells that invaginate to the inside of the embryo can potentially come back out to the surface at a later time, like cells in the cephalic furrow do at ~45 minutes. Since none of our analyses involve "reappearing" cells, in this work we chose to ignore this limitation. However, it is clear that to have a complete representation of all surface cells at later time periods our algorithm will need to be upgraded. One way to do so is by manually marking a watershed seeding point inside each reappearing cell upon its return to the outer surface. Once it is marked, it will continue to be segmented and tracked automatically by the registration-based segmentation mechanism. Another plausible direction, which can be achieved with minor adjustments to the implementation, is to execute our algorithm on the same dataset a second time, after inverting the temporal order of the volumes. When time is reversed, reappearing cells will be segmented and tracked as invaginating cells are in the normal temporal sequence.

### S1. Image analysis methodology.

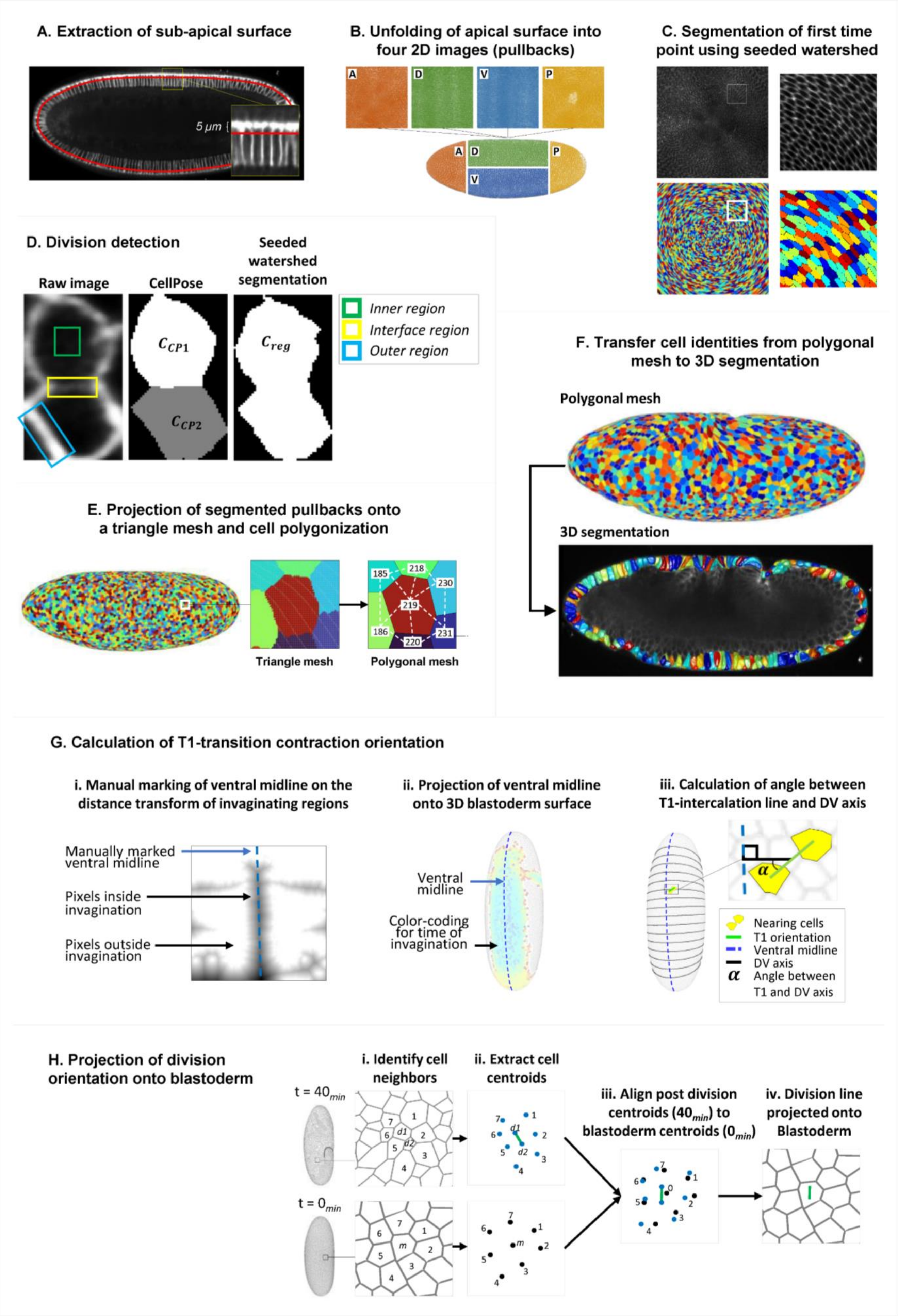

**S1. Image analysis methodology.**

**A-E. Whole embryo single cell segmentation, tracking, division detection and polygonization:** A. A sagittal section of the blastoderm, and a red line marking the surface located 5 microns beneath the apical end of the cells. This surface will be taken for segmentation due to the clear appearance of the membrane. B. Unfolding of the embryo surface into four pullbacks (2D images) corresponding to anterior, posterior, dorsal and ventral regions of the embryo, as calculated by the *ImSAnE* algorithm. Note that to simplify the presentation of this step all anatomical overlaps between pairs of pullbacks have been removed. C. A demonstration of watershed-based segmentation of the anterior pullback of the embryo at the first time point. Top: the raw pullback (left) and a zoomed in box (right), bottom: the same regions as the top after segmentation. Each cell was colored randomly to allow distinguishing between them. D. Detection of divisions in segmented cells. Left: the raw image of a cell that has just completed division. Middle: segmentation by the CellPose algorithm results in the detection of two cells. Right: segmentation of the cell by watershed seed propagation, which does not allow detection of the two cells. Our pipeline decides whether the cell has divided or not based on the geometry of the two presumed daughter cells in the CellPose segmentation (CCP1, CCP2) and based on the ratio of pixel intensities around the daughter cells (see colored rectangles). E. Left and middle: a triangle mesh of the embryo surface 5 microns beneath the apical surface after assigning each triangle the index (color) of the cell to which it belongs. Right: conversion of the triangle mesh into a polygonal mesh. The index of each cell is provided in black over white, and the resulting connectivity graph of the cells is implied by the dashed white lines between their indices.

**F. Segmentation and tracking of cells in 3D:** Top: the resulting polygonal mesh at t=45 minutes (cells are color-coded randomly), bottom: a sagittal slice from the raw volume (gray-scale image in background), overlaid by 3D reconstruction of the segmented cells from the external layer of the embryo. The index of each 3D cell (indicated by cell colors) is copied from the 2D polygon with which it overlaps.

**G. Procedure for calculating T1-transition orientation:** i. We begin by marking the ventral midline of the embryo. To this end, a distance transform map is generated showing the distance of each pixel to the nearest non-invaginating region. Based on the distance patterns the user marks the ventral midline (dashed blue). ii. Ventral view of the embryo showing the ventral midline (dashed blue line) with respect to the invaginating mesoderm (faded blue cells). iii. Calculation of the orientation of a T1-transition with respect to the DV axis of the embryo. Yellow cells – converging cells, green line – the line between the centroids of the converging cells, dashed blue line – the ventral midline as marked by the user, black line – the orientation of the DV axis (orthogonal to the ventral midline), and α – the angle between the T1 orientation line and the DV axis line.

**H. Procedure for projecting division orientations onto blastoderm:** i. The set of all immediate neighbors of the two daughter cells at the first time point following division are identified (top) and are also traced back to the blastoderm (bottom). ii. The centroids of all cells are calculated at both time points. iii. An affine transformation is calculated, which matches the centroids of neighbors after division to the centroids of the same cells in the blastoderm. iv. Using the calculated affine transformation to transfer the division line (green) to the blastoderm.

### S2. Mapping of cell apical area and cell volume ratios.

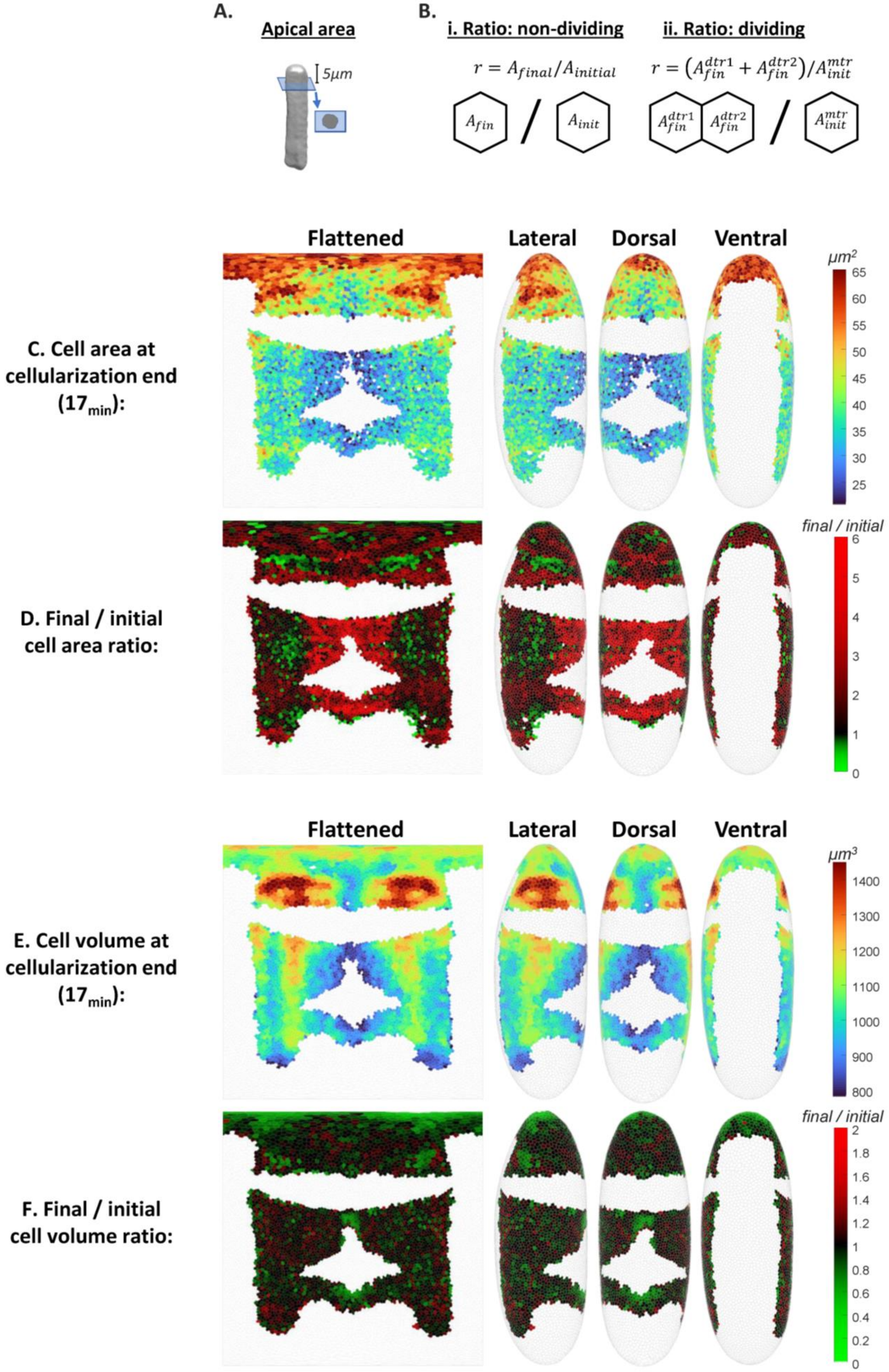

**S2. Mapping of cell apical area and cell volume ratios:**

A. Schematics showing the plane located 5 microns beneath the apical tip of the cell. All calculations of apical area are based on this plane.

B. Schematics of the calculation of apical area ratio in non-dividing (i) and dividing (ii) cells. In dividing cells, the sum of apical areas of both daughter cells is divided by the apical area of the mother cell.

C. Maps showing apical cell areas in non-invaginating cells at $t$=0 minutes.

D. Maps showing apical cell area ratio in non-invaginating cells (values at $t$=43 minutes divided by values at $t$=0 minutes).

E. Maps showing cell volume in non-invaginating cells at $t$=17 minutes, demonstrating complex patterns already at this early stage.

F. Maps showing cell volume ratio in non-invaginating cells (values at $t$=43 minutes divided by values at $t$=0 minutes).

### S3. Mapping of mitotic domains, division times and division orientations.

A. Time of cytokinesis completion

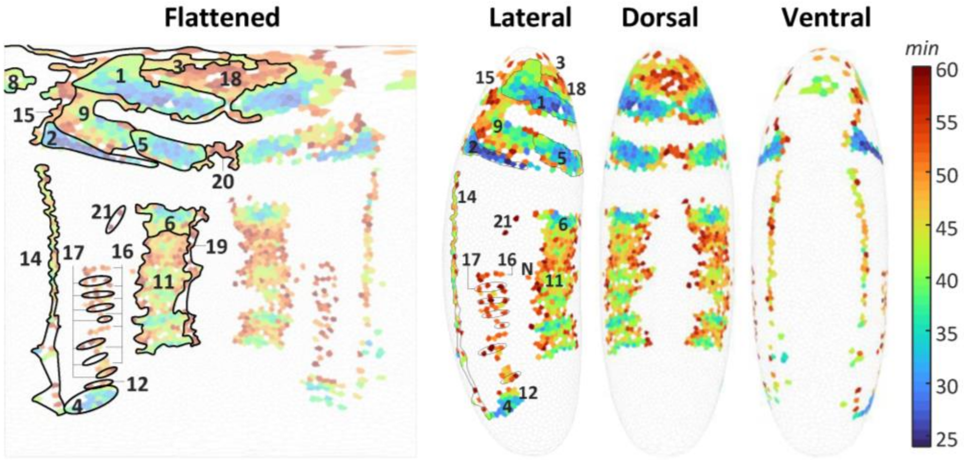


B. The original fate map of mitotic domains proposed by Victoria Foe

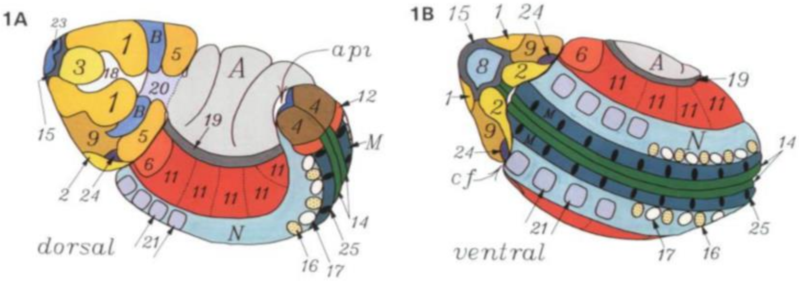


#### C. Mapping of division times times in four additional embryos.

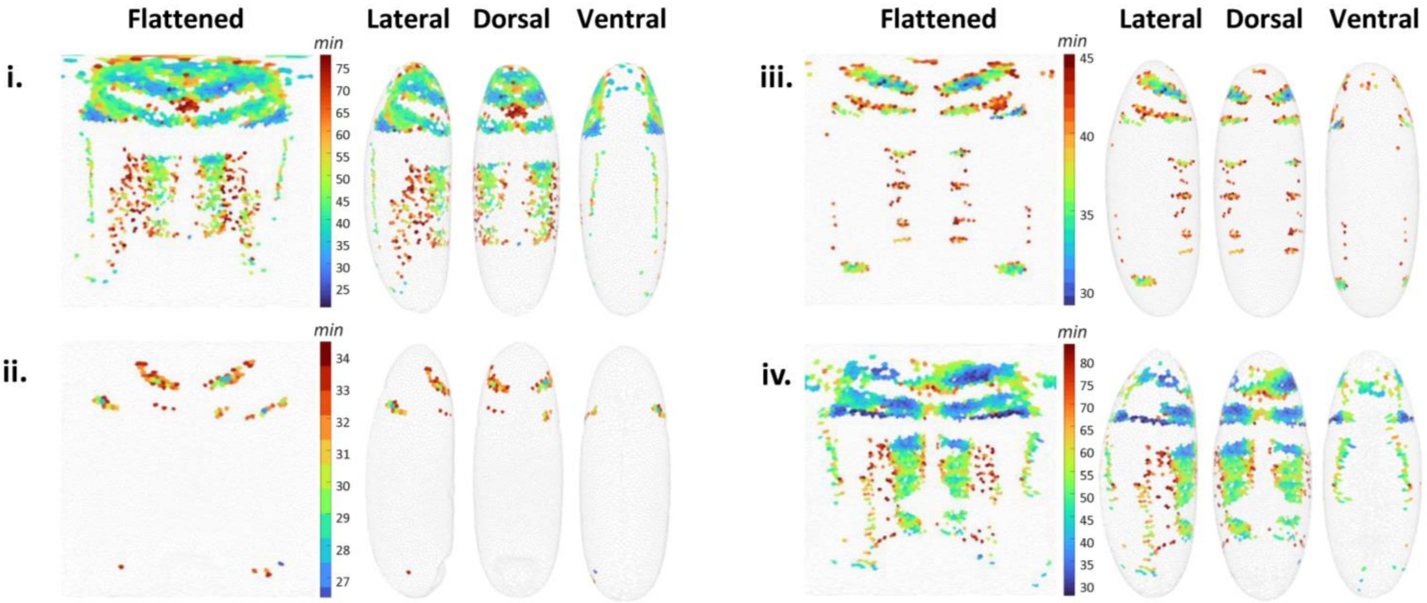

**S3. Mapping of mitotic domains, division times and division orientations.**

A. **Cell division maps.** Each dividing cell is color-coded for the time of cytokinesis completion.

B. **Mapping of mitotic domains, as proposed originally by Victoria Foe:** Comparison of our reconstructed maps with the original maps by Foe shows high agreement in both the anatomical location and timing of the different domain, demonstrating the accuracy of our approach. Figure is copied from: Foe VE. Mitotic domains reveal early commitment of cells in Drosophila embryos. Trends Genet [Internet]. 1989; 5(C):322. Available from: http://linkinghub.elsevier.com/retrieve/pii/0168952589901200.

C. **Four additional examples of mapping cell division events:** Each of the four embryos was imaged over a different period throughout gastrulation, and therefore showing a different subset of mitotic domains.

**Estimated time periods of movies (t=0 is the beginning of apical constriction during mesoderm invagination):**
Embryo in S3A (and in Figure 3A): 5-63 mins.
Embryos S3Ci,ii,iii,iv: 4-77 mins, 21-34 mins, (-5)-45 mins and 14-83 mins, respectively.
* Note that the color-bar associated with each embryo is scaled for the time period in which divisions occur and does not necessarily span over the entire period of the movie.
